# Scopoletin 8-hydroxylase: a novel enzyme involved in coumarin biosynthesis and iron-deficiency responses in Arabidopsis

**DOI:** 10.1101/197806

**Authors:** Joanna Siwinska, Kinga Wcisla, Alexandre Olry, Jeremy Grosjean, Alain Hehn, Frederic Bourgaud, Andrew A. Meharg, Manus Carey, Ewa Lojkowska, Anna Ihnatowicz

**Affiliations:** Intercollegiate Faculty of Biotechnology of University of Gdansk and Medical University of Gdansk, Abrahama 58, 80-307 Gdansk, Poland; Université de Lorraine, UMR 1121 Laboratoire Agronomie et Environnement Nancy-Colmar, 2 avenue de la forêt de Haye 54518 Vandoeuvre-lès-Nancy, France; INRA, UMR 1121 Laboratoire Agronomie et Environnement Nancy-Colmar, 2 avenue de la forêt de Haye 54518 Vandoeuvre-lès-Nancy, France; Institute for Global Food Security, Queen’s University Belfast, David Keir Building, Malone Road, Belfast, UK

**Keywords:** abiotic stress, Arabidopsis, enzyme activity, fraxetin, Fe- and 2OG-dependent dioxygenase, plant–environment interactions, mineral nutrition

## Abstract

**Highlight:** A strongly iron-responsive gene of previously unknown function, At3g12900, encodes a scopoletin 8-hydroxylase involved in coumarin biosynthesis and plays an important role in the iron uptake strategy in Arabidopsis.

**Abstract:** Iron (Fe) deficiency represents a serious agricultural problem, particularly in alkaline soils. Secretion of coumarins by *Arabidopsis thaliana* roots is induced under Fe-deficiency. An essential enzyme for the biosynthesis of major Arabidopsis coumarins, scopoletin and its derivatives, is Feruloyl-CoA 6’-Hydroxylase1 (F6′H1) that belongs to a large enzyme family of the 2-oxoglutarate and Fe(II)-dependent dioxygenases. Another member of this family that is a close homologue of F6’H1 and is encoded by a strongly Fe-responsive gene, At3g12900, is functionally characterized in the presented work. We purified the At3g12900 protein heterologously expressed in *Escherichia coli* and demonstrated that it is involved in the conversion of scopoletin into fraxetin *via* hydroxylation at the C8-position. Consequently, it was named scopoletin 8-hydroxylase (S8H). Its function in plant cells was confirmed by the transient expression of S8H protein in *Nicotiana benthamiana* leaves followed by the metabolite profiling and the biochemical and ionomic characterization of Arabidopsis *s8h* knockout lines grown under various regimes of Fe availability. Our results indicate that S8H is involved in coumarin biosynthesis as part of the Fe acquisition machinery.

## Introduction

Iron (Fe) is an essential micronutrient for all living organisms. In plants, chloroplast and mitochondria have high Fe demand, and play key roles in the Fe homeostasis network in photosynthetic cells (Nouet et al., 2011). Fe is abundant in soils, but its availability is often limited due to soil conditions, which can be highly heterogeneous, such as pH and redox presence of co-elements (Moosavi and Ronaghi, 2011; Marschner, 2012). Fe deficiency is a serious agricultural problem particularly in alkaline and calcareous soils (Mengel, 1994). These types of soils, which represent approximately 30 % of the world’s cropland, are characterized by a higher pH that in combination with the presence of oxygen leads to Fe precipitation in the form of insoluble ferric oxides (Fe_2_O_3_) (Morrissey and Guerinot, 2009).

Higher plants have developed two different types of strategies in response to Fe limitation. In a reduction-based strategy (Strategy I), which occurs in all plant species except grasses, a release of protons into the rhizosphere is enhanced (Marschner and Roemheld, 1994; Kim and Guerinot, 2007) by Fe deficiency-induced proton-translocating adenosine triphosphatases, such as AHA2 in Arabidopsis (Santi and Schmidt, 2009). As a result, at lower pH the ferric oxide precipitates are being dissolved and the plasma membrane bound Ferric Chelate Reductase 2 (FRO2) (Robinson et al., 1999) catalyse the reduction of ferric ions (Fe^3+^) into more soluble and bioavailable to plants ferrous ions (Fe^2+^). Once reduced, Fe^2+^ are transported into the root epidermal cells across the plasma membrane by the divalent metal transporter Iron Regulated Transporter 1 (IRT1) (Vert et al., 2002; Varotto et al., 2002). A second strategy, used by grasses, is based on the release of soluble mugineic acid family phytosiderophores (PS) from the root epidermis and forming the complexes with Fe^3+^ (Takagi et al., 1984; Marschner and Roemheld, 1994; Kim and Guerinot, 2007). The resulting Fe^3+^-PS complexes are then transported into the root epidermis *via* a high-affinity uptake system without the requirement of a reduction step (Curie et al., 2001; Kim and Guerinot, 2007).

The precise mechanisms underlying responses of Strategy I plants to low Fe availability in calcareous soils are not clear. However, it is well documented that Fe deficiency enhanced release of reductants/chelators (mainly phenolics) in many dicots (Marschner and Roemheld, 1994; Jin et al., 2007). Recently, it was shown that Fe deficiency induces the secretion of secondary metabolites like scopoletin and its derivatives by Arabidopsis roots (Rodriguez-Celma et al., 2013a; Fourcroy et al., 2014), and that Feruloyl-CoA 6’-Hydroxylase1 (F6’H1) is required for the biosynthesis of the Fe(III)-chelating coumarin esculetin which is released into the rhizosphere as part of the Strategy I-type Fe acquisition machinery (Schmid et al., 2014).

Scopoletin and, its corresponding glycoside, scopolin are the predominant coumarins in Arabidopsis roots. But simultaneously, many other coumarin compounds such as skimmin, esculetin, fraxetin and recently discovered coumarinolignans are present in the roots of this model plant (Rohde et al., 2004; Bednarek et al., 2005; Kai et al., 2006; Kai et al., 2008; Schmid et al., 2014; Schmidt et al., 2014; Ziegler et al., 2016; Siso-Terraza et al. 2016; Ziegler et al., 2017). In our previous work, we reported the presence of a significant natural variation in scopoletin and scopolin accumulation between various Arabidopsis accessions (Siwinska et al., 2014). It is an interesting issue due to the fact that Strategy I plants differ considerably between plant species and genotypes in their tolerance to Fe deficiency. Among coumarins significantly highly accumulated in response to Fe limited condition, scopoletin and fraxetin together with their corresponding glycosides were detected (Fourcroy et al., 2014; Schmid et al., 2014; Schmidt et al., 2014). Up to now no enzymes involved in the last step of fraxetin biosynthesis were identified.

The above findings point out that the biosynthesis of coumarins and their accumulation are related to plant responses to Fe deficiency, but the exact mechanisms of action underlying these processes have remained largely unknown. To better understand these mechanisms, we selected and functionally characterize an enzyme of unknown biological role encoded by the At3g12900 gene, which share significant homologies with the F6’H1 and F6’H2 described by Kai et al. (2008) as involved in the synthesis of scopoletin, and which is pointed in the literature as a strongly iron-responsive (Lan et al., 2011; Rodriguez-Celma et al., 2013b; Mai et al., 2015; Mai and Bauer, 2016). This work lead us to identify the At3g12900 oxidoreductase as a scopoletin 8-hydroxylase (S8H) involved in the biosynthesis of fraxetin that is associated with regulation of Fe homeostasis in Arabidopsis.

## Materials and methods

### Plant material

*Arabidopsis thaliana* (Arabidopsis) Col-0 accession was used as the wild-type. The *s8h-1* (SM_3.27151) and *s8h-2* (SM_3.23443) T-DNA insertional mutant lines (Figure S1) were identified in the SM collection (http://signal.salk.edu/; Tissier et al., 1999) of single copy dSpm insertions (Col-0 background). Seeds of both *s8h* lines are available at the stock centre NASC (http://arabidopsis.info/). Genotyping of *s8h-1* and *s8h-2* was done using protocols and primers described in Table S1. Selected *s8h-1* homozygous mutants were tested for *S8H* gene expression by qRT-PCR (Figure S2, Table S2). A lack of *S8H* transcript in *s8h-2* mutant background was confirmed by RT-PCR using primers shown in Table S2. *Nicotiana benthamiana* seeds were kindly gifted by Dr Etienne Herbach (INRA, Colmar, France).

### Growth conditions

#### Hydroponic cultures

Plants were grown in two types of modified Heeg solutions (Heeg et al. 2008) described in details in Table S3. First solution (10xHeeg) was prepared as previously described (Ihnatowicz et al., 2014), a second solution (1xHeeg) contained 10 times less microelements (except for Fe^2+^). Arabidopsis seeds were surface sterilized by soaking in 70 % ethanol for 2 min and subsequently kept in 5 % calcium hypochlorite solution for 8 min. Afterwards seeds were rinsed three times in autoclaved deionized water. The surface-sterilized seeds were sown out ton tip-cut 0.65% agar-filled tubes or on micro-centrifuge tubes’ lids (filled with solidified Heeg medium) that were placed into tip boxes with control hydroponic solution (40 μM Fe^2+^). After a few-days stratification at 4 °C, boxes with plants were kept in controlled environment (16 h light at 22 °C/∼7000 lux and 8 h dark at 20 °C). Approximately three weeks later, plants were transferred either to a freshly made control solution or to Fe-deficient (10 μM Fe^2+^) or Fe-depleted (0 μM Fe^2+^) solutions. In the first sets of experiments hydroponic solutions were fully changed once per week, while in the second sets of experiments boxes containing old nutrient solution were refilled by the addition of a fresh medium.

#### In vitro cultures on plates

The surface-sterilized seeds were sown out on Petri dishes containing different homemade Murashige and Skoog’s (MS) medium: (1) half strength MS medium (0.5 MS) with macro-elements in half strength and micro-elements/vitamins in full strength and (2) one fourth strength MS medium (0.25 MS) with macro-elements in one fourth strength and microelements/vitamins in half strength. Both type of media contained 1% sucrose, 0.8 % agar supplemented with 4 mg/l glycine, 200 mg/l myo-inositol, 1 mg/l thiamine hydrochloride, 0.5 mg/l pyridoxine hydrochloride and 0.5 mg/l nicotinic acid. For stratification, plates were kept in the dark at 4 °C for 72 h and then placed under defined growth conditions (16 h light (∼5000 lux) at 22 °C and 8 h dark at 20 °C).

#### In vitro liquid cultures

10 days old seedlings from agar plates were transferred into 250 ml glass containing 20 ml sterile 0.25 MS liquid medium. Plants grown in liquid cultures were incubated on rotary platform shakers at 120 rpm. 10 days after transfer, 20 ml of fresh medium was added to each flask. 18 days after transfer, plants were harvested (29th day of culture), leaves and roots were frozen separately in liquid nitrogen and stored at −80 °C.

Two weeks after germination, *N. benthamiana* seeds were transplanted and cultured independently for 3 additional weeks in plant growth chambers under a photoperiod of 16 h light (120 μmol m^−2^ s^−1^) at 24 °C and 8 h dark at 22 °C with 70 % humidity.

### Chemicals

Acids: ferulic (Aldrich), coumaric (Sigma), cinnamic (Sigma) and caffeic (Fluka). Coumarins: coumarin (Sigma), daphnetin (Sigma), esculetin (Sigma), esculin (Sigma), fraxetin (Extrasynthèse), fraxin (Extrasynthèse), isoscopoletin (Extrasynthèse), limetin (Herboreal), scoparon (Herboreal), 6-methoxycoumarin (Apin Chemicals), 7-methoxycoumarin (Herboreal), scopoletin (Herboreal), scopolin (Aktin Chemicals Inc.), umbelliferon (Extrasynthèse), skimmin (Aktin Chemicals Inc.) and 4-methylumbelliferon (Sigma). The CoA thiol esters of the cinnamates (cinnamoyl CoA, *p*-coumaroyl CoA, caffeoyl CoA and feruloyl CoA) were enzymatically synthetized as described by Vialart et al. (2012). *P*-coumarate and coenzymeA (CoA) were purchased from Sigma-Aldrich. Kanamycin, chloramphenicol and isopropyl-β-d-thio-galactopyrannoside (IPTG) were purchased from Duchefa.

### Construction of binary vector and Agrobacterium tumefaciens strains

The amplified *S8H* ORF (details of PCR in Table S4) was first cloned into the pCR8 plasmid using the pCR®8/GW/TOPO® TA cloning kit (Invitrogen) (Vialart et al., 2012) using the Gateway technology. The recombinant pBIN-GW-S8H vector was further introduced into the LBA4404 *A. tumefaciens* strain and used together with the C5851 *A. tumefaciens* strain containing pBIN61-P19 (Voinnet et al., 2003, provided by D. Baulcombe (Department of Plant Science, University of Cambridge, UK)) for transient expression in *N. benthamiana* leaves.

### Construction of pET28a+ expression vector and E. coli Rosetta 2 strain

The ORF of *S8H* was amplified by PCR (5’ primer: 5’-<u>GGATCC</u>GGTATCAATTTCGAGGACCAAAC-3’ and 3’ primer: 5’-<u>CTCGAG</u>CTCGGCACGTGCGAAGTCGAG-3’) and cloned between the *Bam*HI and *Xho*I sites of pET28a+. The recombinant plasmid was introduced into competent *E. coli* Rosetta 2 (Novagen) strain by heat shock.

### Heterologous expression and purification of S8H

*E. coli* Rosetta 2 strain transformed with pET28a+-S8H was cultured at 37 °C overnight in 10 ml LB (Sambrook, 2001) supplemented with 100 mg/L kanamycin and 33 mg/L chloramphenicol. A 2 ml preculture was transferred to 1L of fresh LB containing kanamycin 100 mg/L and chloramphenicol 33 mg/L. Induction of the S8H expression was adapted from the protocol developed by Oganesyan et al. (2007). Transformed cells were cultured at 37 °C until OD_600nm_ reached 0.6. A salt stress with 0.5 M NaCl and heat stress at 47 °C were applied during 1 h in the presence of 2 mM betain. Then temperature was set at 20 °C for 1 h and finally the expression of S8H was initiated by adding 1 mM of IPTG. Cells were harvested after 14 h by a 20 min centrifugation at 4000 g at 4 °C and the pellet was resuspended in 4 ml of potassium phosphate buffer pH 8.0 with 10 mM imidazol. The cell suspension was sonicated (Bandelin SONOPLUS apparatus; for 20 sec x 5 with an interval of 50%, 60% power) and subsequently centrifuged at 10000 g for 20 min at 4 °C. The purification of soluble recombinant his-tagged proteins was done using the TALON Metal Affinity Resin (TAKARA) as described by the supplier. The purified proteins were eluted with 0.1 M potassium phosphate buffer, 200 mM imidazol solution (pH 8.0).

### Enzymatic activities measurements

Enzymatic incubations were made immediately after the S8H purification. The protocol was adapted from Kai et al. (2008). The reaction was done in 200 μl volume at saturating concentration of FeSO_4_ (0.5 mM), α-cetoglutarate (5 mM), sodium ascorbate (5 mM) in 0.1 M potassium phosphate buffer at optimal pH (7.0), 200 μM substrate and 2.6 μg of purified enzyme. Reaction mixtures were incubated for 10 min at optimal temperature of 31.5 °C. Enzymatic incubations with CoA esters were additionally incubated at the end of the reaction with 20 μl of 5 M NaOH for 20 min at 37 °C to hydrolase the ester bond and subsequently with 20 μl of acetic acid in order to close the lacton ring. Samples with acids and coumarins as substrates were stopped by the addition of 2 μl of trifluoroacetic acid and incubation in 20°C for 20 min. Reaction mixtures were then centrifuged for 30 min at 13000 rpm, the resulting supernatant was recovered, filtrated (0.22 μm) and analysed by HPLC.

### High Performance Liquid Chromatography (HPLC)

The separation was performed using HPLC Shimadzu system with DAD/UV detector. Injected volume was 50 μl. Samples were separated on Interchim C18 Lichrospher OD2(250*4.0 mm, 5 μm) column with the flow speed 0.8 ml/min. Separation was performed by elution with following program (A) H_2_O and 0.1 % acetic acid (B) methanol and 0.1 % acetic acid: 0-35 min gradient 10 % - 70 % B, 35-36 min 70 % - 99 % B, 36-39 min 99 % - 99 % B, isocratic elution and column regeneration. Wavelength was set at 338 nm to monitor the formation of fraxetin.

### UHPLC and LTQ LC/MS analysis

Sample analysis were performed on a Nexera UHPLC system (Shimadzu Corp., Kyoto, Japan) coupled with LCMS 2020 mass spectrometer (Shimadzu). Chromatographic column was a C18 reverse phase (Zorbax Eclipse Plus, Agilent technologies, Santa Clara, CA, USA) 150*2.1mm 1,8μm. Elution of the compounds was performed as described in Dugrand et al. (2013). To confirm the *in vitro* metabolisation of scopoletin into fraxetin, separation were processed on a Ultimate 3000 chromatographic chain coupled with a LTQ-XL mass spectrometer (Thermo Electron Corporation, Waltham, MA, USA) as described by Karamat et al. (2012).

### Heterologous expression in N. benthamiana leaves

The protocol was adapted from Voinnet et al. (2003). Several freshly spread colonies (LBA4040 [pBIN-F6’H2], LBA4040 [pBIN-S8H)], C5851 [pBIN61-P10]) were inoculated to 40 ml LB medium containing the proper antibiotics and incubated at 28 °C for one night. The cultures were centrifuged for 10 min at 4000 g and the pellet resuspended in 20 ml sterile deionizated water. This step was repeated 3 times in order to remove all traces of antibiotics. *N. benthamiana* leaves were infiltrated with recombinant strains in order to have an OD_600_ of 0.4 for pBIN61-P19 bacteria and 0.2 for pBIN-S8H or pBIN-F6’H2 bacteria. The infiltrated plants were stored in a growth chamber during 96 h. In parallel, for each experiment, *N. benthamiana* were infiltrated with LBA4404 [pBIN-GFP] as a positive control. Expression of GFP was checked under a binocular microscope.

### *Extraction of polyphenols from N. benthamiana* leaves

Freshly harvested infiltrated *N. benthamiana* leaves were crushed in liquid nitrogen and 100 mg mixed to 800 μl 80 % methanol. This solution was vigorously mixed during 30 sec prior a 30 min centrifugation at 10000 g. The supernatant was transferred to a fresh tube and the pellet submitted to a second extraction with 800 μl 80 % methanol. Both supernatants were pooled and vacuum dried. The pellet was resuspended in 100 μl MeOH/H2O 80/20 v/v and analyzed by UHPLC.

### Preparation of methanol extracts from Arabidopsis roots

Arabidopsis roots were frozen in liquid nitrogen and grinded with pestle and mortar. 50 mg of plant tissue were mixed in 500 μl of 80 % methanol supplemented with 2.5 μM 4-methylumbelliferon as an internal standard. Samples were sonicated for 10 min and centrifuged for 10 min at 10 000 g. Supernatants were transferred to fresh tubes, vacuum dried, resuspended in 100 μl MeOH/H_2_O 80/20 v/v and analyzed by UHPLC.

### Extraction of root exudates from nutrient solutions

Nutrient solutions of Arabidopsis *in vitro* cultures were collected 18 days after the onset of Fe treatments. Phenolic compounds were retained in a BAKERBOND™ C18 column (J. T. Baker Chemical Co., Phillipsburg), eluted from the cartridge with 3 ml of 100 % methanol and dried in centrifugal evaporator. Dry extracts were stored at −20 °C for further analysis.

### Trace element analysis

Arabidopsis roots were first lyophilized and subsequently milled before microwave (MARS) assisted digestion in concentrated nitric acid (Aristar). Metal concentrations were determined by Inductively Coupled Plasma-Mass Spectrometry (ICP-MS) (Thermo iCAP Q, Bremen, Germany).

### Chlorophyll measurement

Chlorophyll extraction with acetone was done by the method given by Porra et al. (1989). The absorbances of the diluted supernatants were taken at 750.0, 663.6 and 646.6 nm. After measurement a special formula was used to convert absorbance measurements to mg of Chl (Porra et al., 1989).

### Statistical analysis

All treatments included at least three biological replicates. Data processing and statistical analyses-pairwise comparisons using t-tests (except trace element analysis) were carried out using Excel (Microsoft Excel 2010). Error bars representing standard deviation (SD) are shown in the figures; the data presented are means. The significance level used here is p <0.01 indicated by (**) or p < 0.05 indicated by (*). Statistical significance of differences observed in trace element analysis were analyzed by R version 3.3.2 (2016-10-31) (R Core Team 2014) with the use of agricolae package; p < 0.05 was used. Values that are significantly different are indicated by different letters.

## Results

### Phylogenetic analysis of genes encoding Fe(II)- and 2OG-dependent dioxygenases from Arabidopsis genome

More than 100 genes encoding enzymes sharing sequence homologies with dioxygenases were identified in the genome of Arabidopsis (Kawai et al., 2014). Almost half of them display characteristic amino acid sequence motifs involved in binding cofactors such as Fe^2+^ and 2OG (His-X-Asp-X-His and Arg-X-Ser, respectively; Wilmouth et al., 2002). We performed phylogenetic analysis based on the nucleic sequences of putative genes encoding dioxygenases collected from TAIR. This analysis pointed out that two genes involved in scopoletin biosynthesis (At3g13610 and At1g55290, encoding F6’H1 and F6’H2 respectively) were clustered together and identified the At3g12900 gene of unknown function to share the highest homology with both scopoletin synthases (Figure 1).

**Figure 1.**
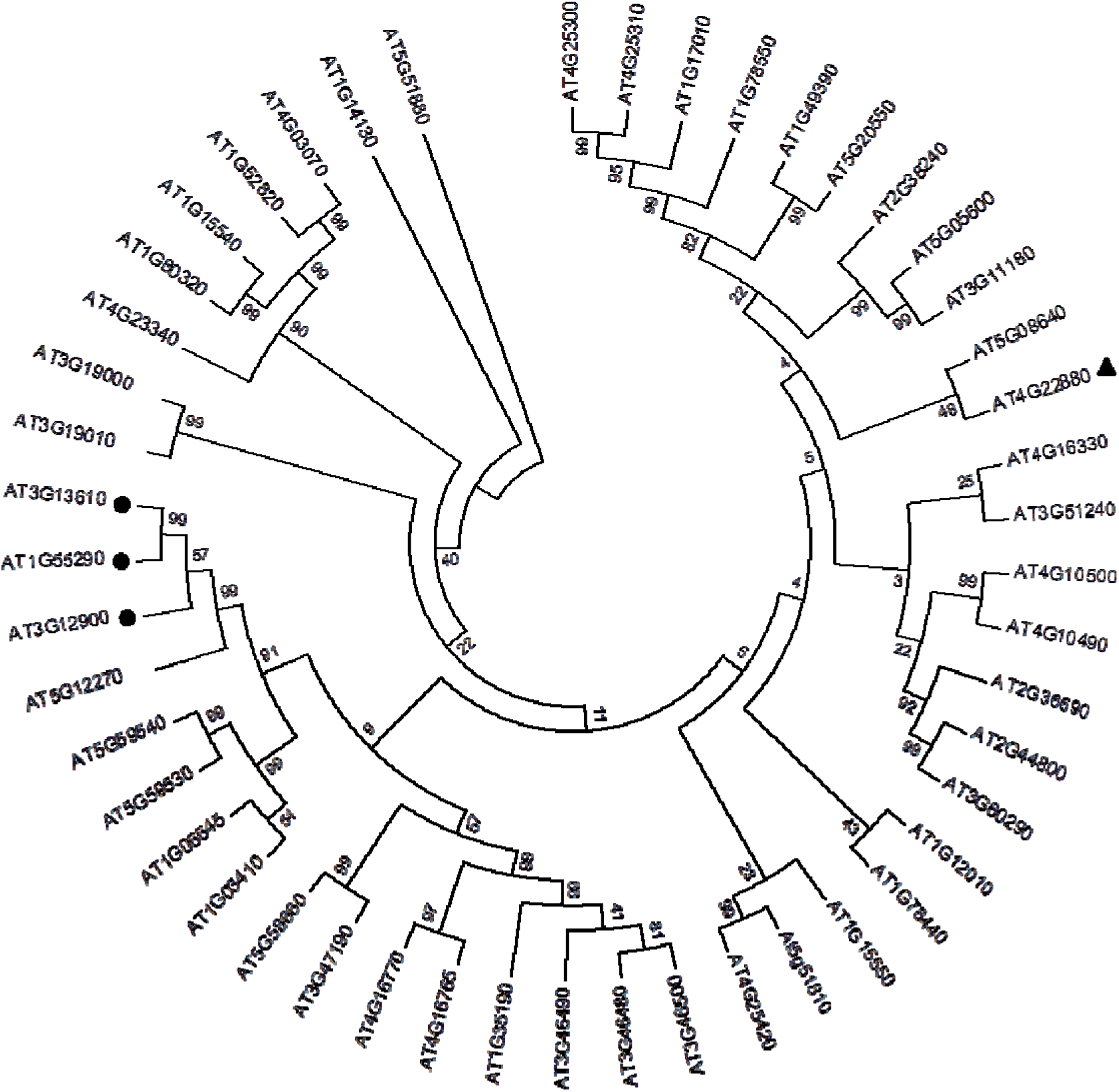
Nucleic sequence based tree. The phylogenetic analysis was performed on available nucleic sequences of genes encoding 2-oxoglutarate (2OG)- and ferrous iron Fe(II)-dependent dioxygenase (2OGD) from Arabidopsis genome. The phylogenetic tree was done using the MEGA 6 software. Black dots (•) indicate genes (At3g13610 and At1g55290) encoding enzymes involved in scopoletin biosynthesis (F6’H1 and F6’H2 respectively), and their closest homologue based on the nucleic sequences - a gene (At3g12900) encoding a dioxygenase of unknown biological function (characterized in this study as S8H). Black triangle (▴) indicate gene (At4g22880) encoding anthocyanidin synthase (ANS) enzyme. Nucleic sequences downloaded from TAIR (http://arabidopsis.org/) were selected based on the presence of sequences coding for Fe (His-X-Asp-X-His) and 2OG (Arg-X-Ser) binding motifs (Wilmouth et al. 2002).

The amino acid sequences of three enzymes were compared to a leucoanthocyanidin dioxygenase (named anthocyanidin synthase - ANS) encoded by At4g22880 which catalyses the conversion of leucoanthocyanidins into anthocyanidins (Heller and Forkmann, 1986) and has the highest sequence identity to F6’H1 among enzymes of known crystallographic structure (Wilmouth et al., 2002). The Fe^2+^ binding site (His 235, His 293, Asp 237 in F6’H1) and the 2OG binding site (Arg 303 and Ser 305 in F6’H1) are highly conserved in all proteins (Figure S3). In contrast, the amino acid residues responsible for the substrates binding differ among the enzymes which is consistent with the activities already described for three of them: feruloyl-CoA ester for F6’H1 and F6’H2 (Kai et al., 2008) and leucoanthocyanidin for ANS (Wilmouth et al., 2002). These alignments highlight a different sequence for At3g12900 which suggest another substrate specificity.

### Determination of in vitro substrate specificity and kinetic parameters of enzymatic reaction catalysed by the At3g12900 oxidoreductase

In order to determine the substrate specificity, the At3g12900 6XHis-tagged protein was expressed in *Escherichia coli*. An improved protocol was used for the purification of heterologously expressed S8H enzyme to obtain the purified protein fraction (Figure S4). The purified enzyme was incubated in the presence of dioxygenase cofactors at saturating concentrations and 19 various potential substrates belonging to (1) cinnamoyl derivative CoA esters; (2) cinnamic derivative acids; (3) and coumarins (Table S5). Only scopoletin was converted into a product (Figure 2A) which has been unambiguously identified as fraxetin due to its UV absorption spectrum, its molecular mass (Figure 2ABC) and its MS fragmentation spectrum in comparison to a fraxetin commercial standard (Figure 2DE). The At3g12900 can therefore be considered as a scopoletin 8-hydroxylase (S8H) (Figure 3) catalysing the hydroxylation at the position C8 of scopoletin. The optimal reaction conditions were determined to be at pH 8.0±0.1 at 31.5±0.4 °C (Figure 4AB) and the optimal incubation time was set at 10 minutes (Figure S5A). These experimental conditions led us to determine the kinetic characteristics of the enzyme as being *K*_*m*_ 11±2 μM, V_max_ 1.73±0.09 pmol/sec/pmol of S8H (Figure S6). We also demonstrated that the enzyme efficiency is the best in presence of 50 μM Fe^2+^ (Figure S5B) and that higher concentrations significantly reduced the enzyme activity leading to a decreased fraxetin synthesis.

**Figure 2.**
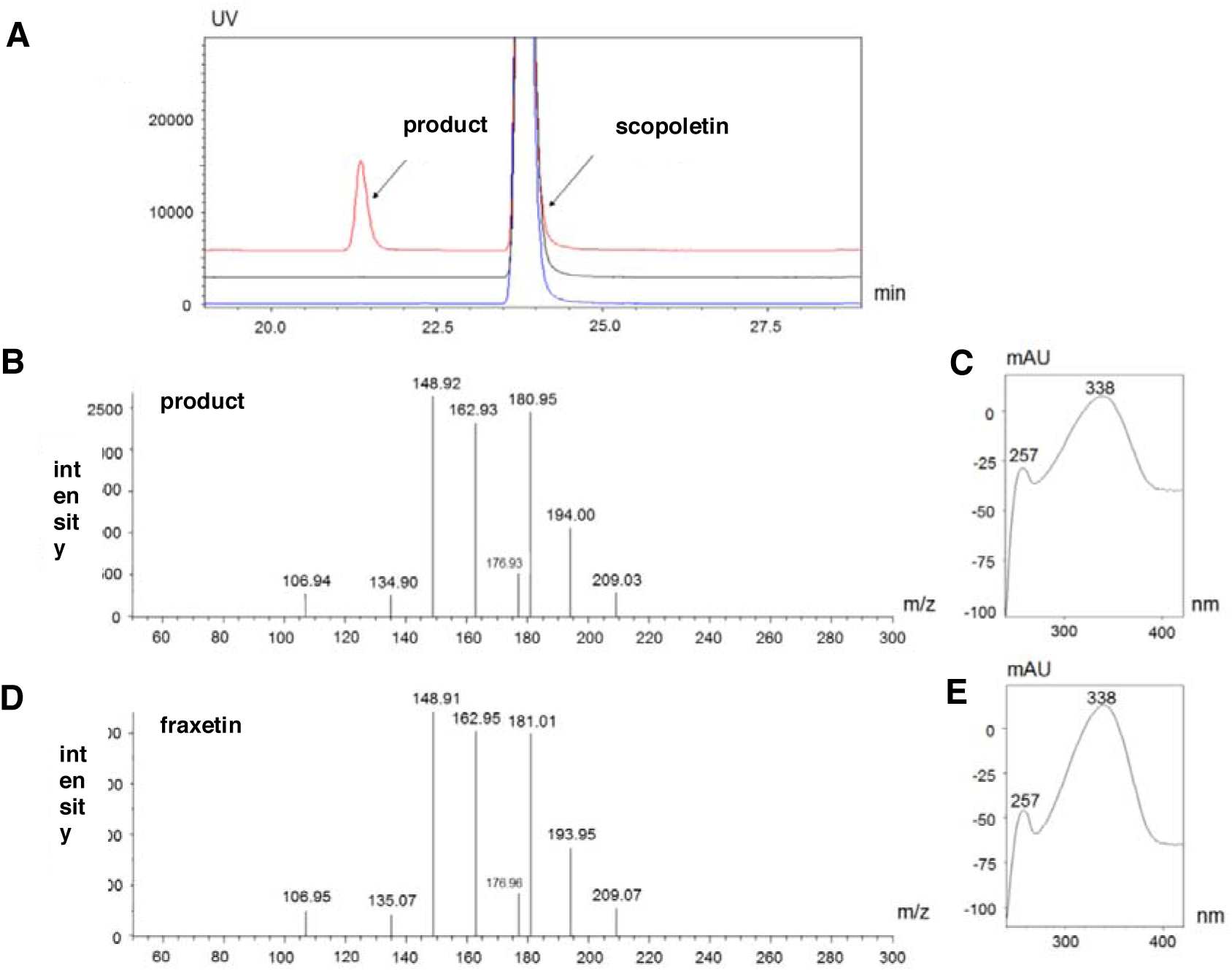
*In vitro* functional characterization of the S8H enzyme. **(A)** The chromatograms in blue and black are the two negative control reactions. Blue show the reaction carried out without the addition of iron, and black without the addition of oxoglutarate. The chromatogram in red shows the result of S8H enzyme incubation with scopoletin and enzyme cofactors – Fe(II) and 2OG. **(B)** MS fragmentation of the additional peak from the chromatogram shown in red in point A. **(C)** UV absorbance spectrum of the additional peak from the chromatogram indicated in red. **(D)** MS fragmentation of fraxetin standard. **(E)** UV absorbance spectrum of fraxetin standard.

**Figure 3.**
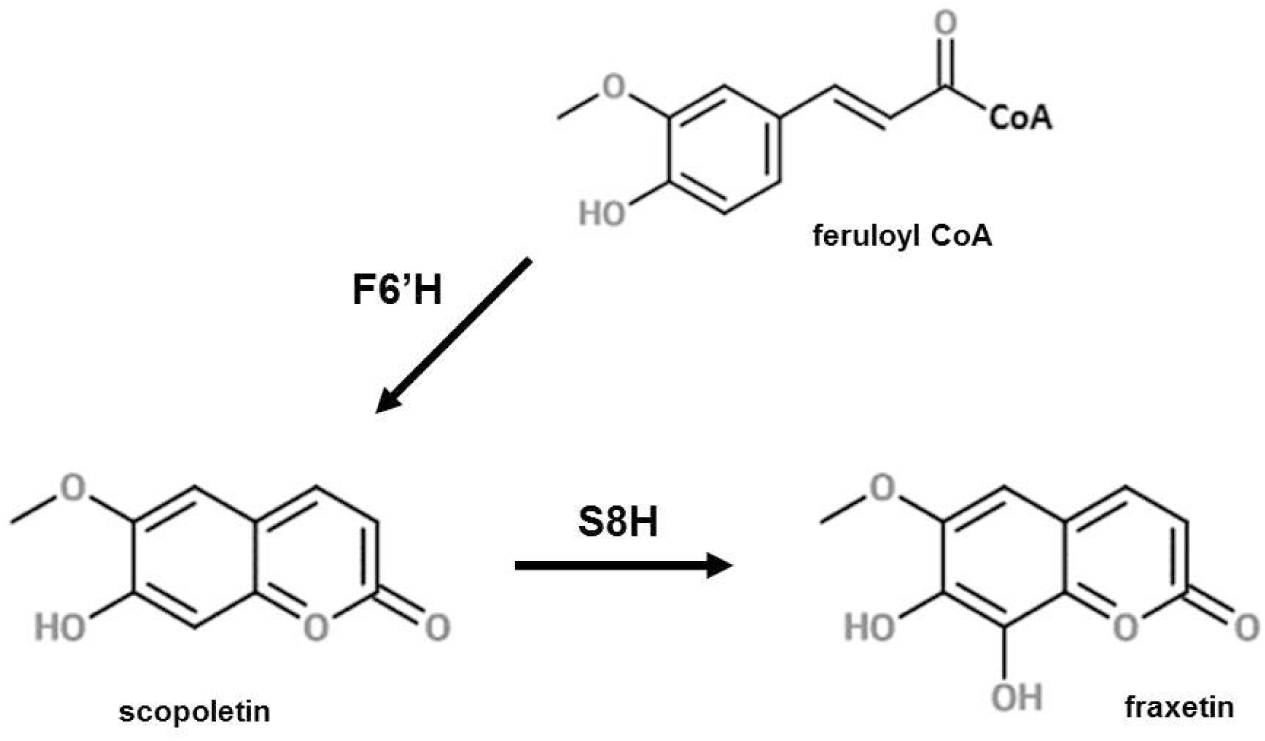
Last step of fraxetin biosynthesis in *Arabidopsis thaliana* catalyzed by scopoletin 8-hydroxylase (S8H). The Fe(II)- and 2-oxoglutarate-dependent dioxygenase (F6’H) catalyses the ortho-hydroxylation of feruloyl CoA before the lactone ring formation of scopoletin. Subsequently, S8H catalyses the hydroxylation at the position C8 of scopoletin leading to fraxetin production.

**Figure 4.**
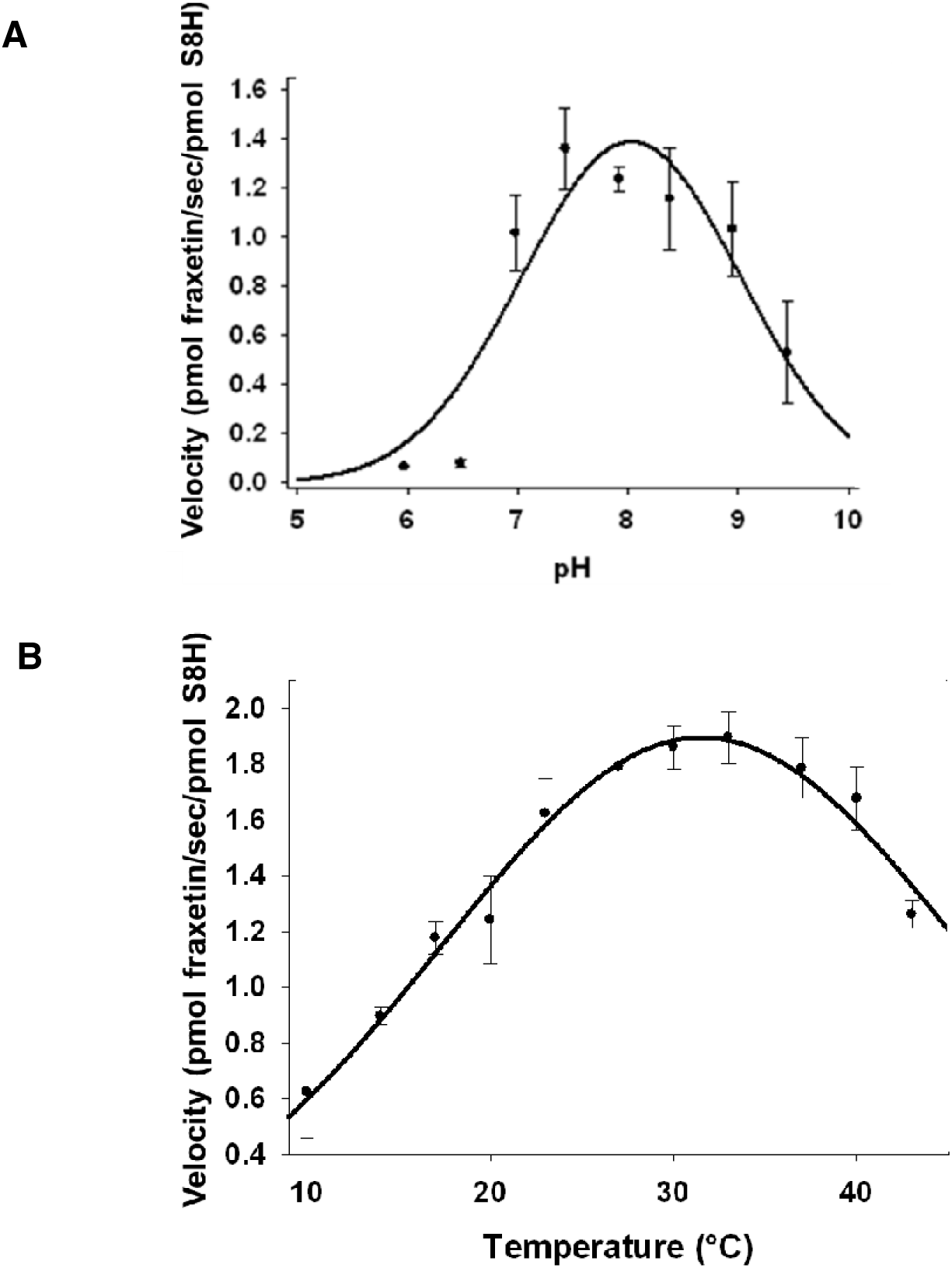
The optimal pH and temperature for *in vitro* activity of S8H. **(A)** pH dependent activity of *Arabidopsis thaliana* S8H. The reactions were performed in triplicate at optimal temperature of 31.5 °C at 200 μM of scopoletin, 5 mM of alpha-acetoglutarate and 0.2 mM FeSO_4_. pH range was 5.02-9.45 in buffer Tris 0.1 M. Reactions were monitored at 340 nm and scopoletin product was quantified with a standard curve. Nonlinear regression analysis against equation f = a*exp(-0.5*[(x-x_0_)/b)^2^] was performed with Sigma Plot 12.0 Software. Optimal pH was 8.0 +/- 0.1. **(B)** Temperature dependent activity of Arabidopsis S8H. The reactions were performed in triplicate at optimal pH of 8.0 at 200 μM of scopoletin, 5 mM of alpha-acetoglutarate and 0.2 mM FeSO_4_. Temperature range was 10-43 °C in buffer Tris 0.1 M. Reactions were monitored at 338 nm and fraxetin product was quantified with a standard curve. Nonlinear regression analysis against equation f = a*exp(-0.5*[(x-x_0_)/b)^2^] was performed with Sigma Plot 12.0 Software. Optimal temperature was 31.5 +/- 0.4 °C.

### In vivo activity of scopoletin 8-hydroxylase in Nicotiana benthamiana

In order to confirm its function in plant cells, we transiently expressed the S8H enzyme in *N. benthamiana* leaves. Our analysis showed that the presence of this enzyme did not increase the production of fraxetin in comparison to a control infiltration performed with bacteria transformed with an empty vector (Figure 5). However, when we conducted a double infiltration leading to the overexpression of S8H and F6’H in tobacco leaves, we detected a significantly higher level of fraxetin in comparison to leaves transiently expressing S8H alone (p = 0.01) or leaves infiltrated with control (the empty vector, p = 0.07) (Figure 5). This result is consistent with an increase of the scopoletin pool that occurs due to the F6’H activity, which can be further transformed into fraxetin and confirms that At3g12900 functions as a scopoletin 8-hydroxylase.

**Figure 5.**
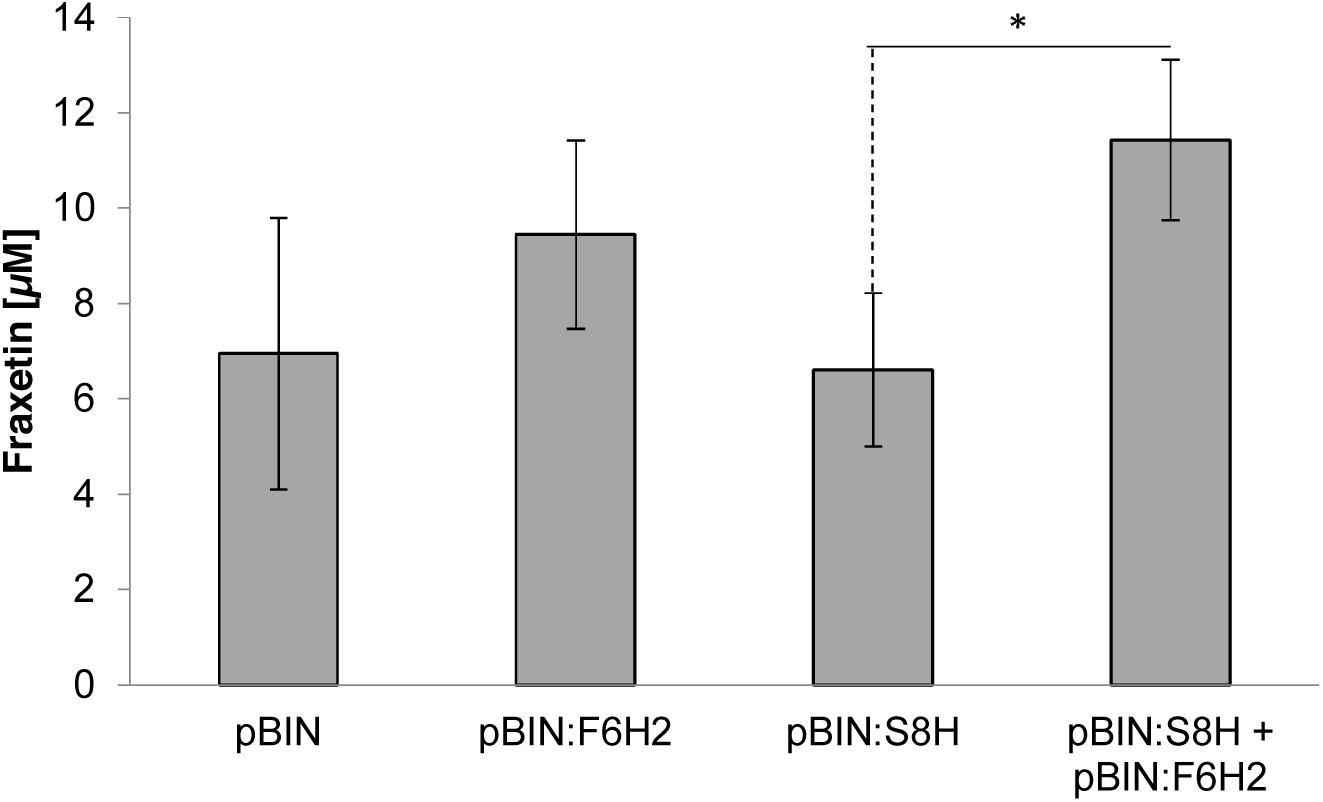
Fraxetin content in *N. benthamiana* leaves. Transient expression was carried out using *A. tumefaciens* with the pBIN empty vector (control), pBIN:S8H, pBIN:F6’H2 and simultaneously with vectors encoding F6’H2 and S8H. Fraxetin content was quantified with UHPLC. (*) p < 0.05.

### Characterization of independent s8h mutant alleles grown in Fe-depleted hydroponics

To get insight into the physiological role of S8H, we identified two independent mutant lines with non-functional S8H dioxygenase (*s8h-1* and *s8h-2*) and investigated their behaviour in various types of culture. Since the literature data reported that fraxetin accumulation is induced under Fe deficiency, firstly we investigated their behaviour in controlled Fe-depleted hydroponic solution. Plants were grown in a control hydroponic solution (40 μM Fe^2+^) for three weeks and subsequently were transferred to Fe-depleted solution (0 μM Fe^2+^) and cultured for additional three weeks. In Fe-depleted solutions both mutant lines were clearly paler than Col-0 control plants (Figure 6A). The mutant phenotype was linked with lower chlorophyll a+b content and chlorophyll a/b ratio (Figure 6B). The targeted metabolite profiling of methanolic extracts from the plant roots grown in hydroponic systems showed a highly significant increase in scopolin (P < 0.01) and scopoletin (P < 0.01) concentration in roots grown in Fe-depleted conditions for both mutant lines and a significant increase of umbelliferon accumulation in *s8h-1* compared with Col-0 plants (Figure 6C).

**Figure 6.**
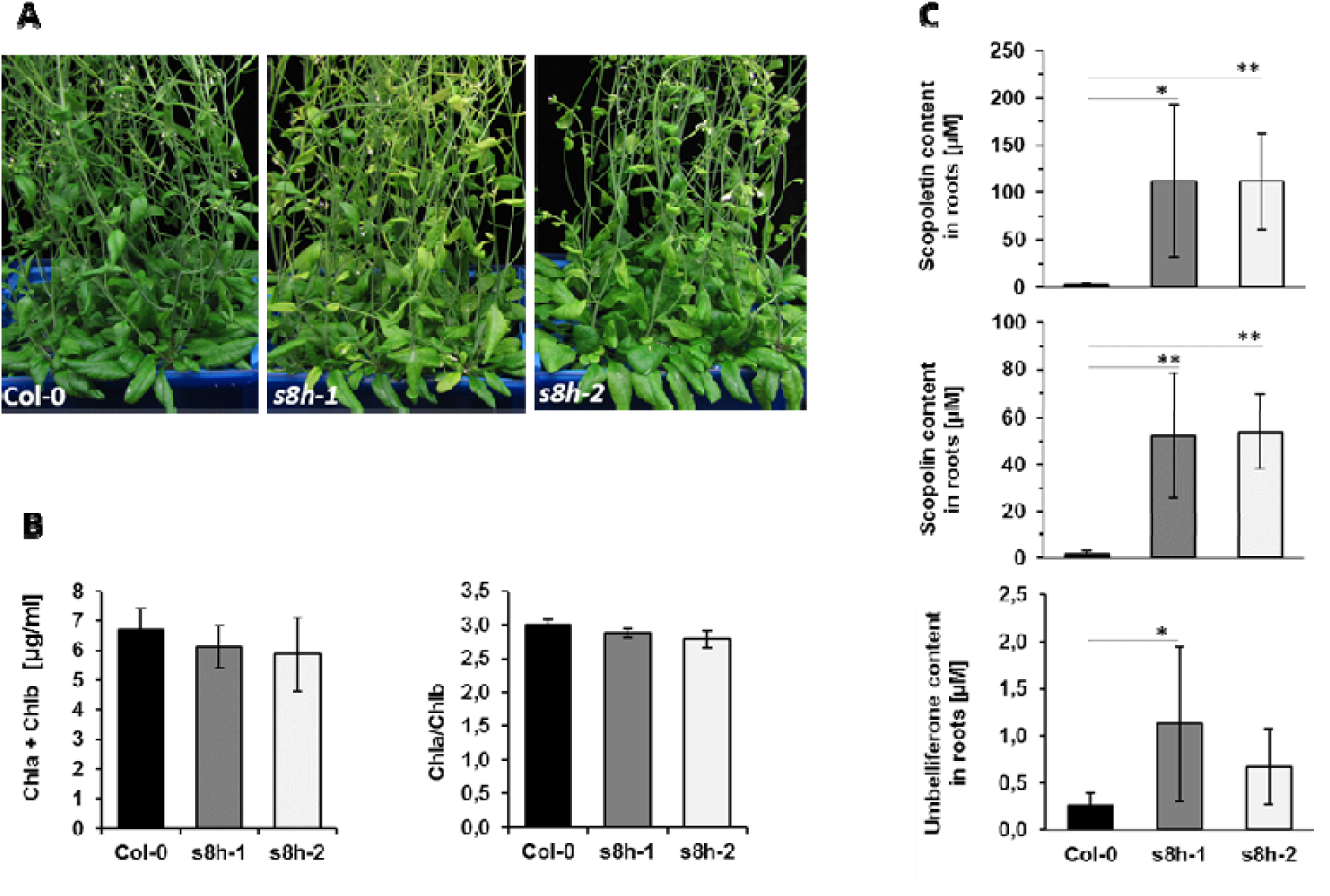
Phenotypic characterization of 6-weeks-old Col-0 plants and two *s8h* T-DNA mutant lines (*s8h-1* and *s8h-2*) grown in Fe-depleted hydroponic solution (0 μM Fe^2+^). After three weeks of growth in optimal hydroponic solution (40 μM Fe^2+^) plants were transferred to **(A)** a freshly made Fe-depleted solutions and cultured until the chlorotic phenotype was clearly visible. Hydroponic solutions were fully changed once per week. **(B)** Chlorophyll concentration and ratio of WT and mutant plants. **(C)** Relative levels of scopoletin, scopolin and umbelliferon accumulated in the plant roots. Metabolite profiling of coumarins were done by UHPLC. The results of one representative experimental replicate are presented. Error bars represent the SD from six biological replicates. (*) p < 0.05, (**) p < 0.01.

### Biochemical and ionomic characterization of plants cultivated in liquid cultures with various Fe content

Since coumarins are mostly stored in roots, all tested genotypes were cultivated *in vitro* in liquid cultures in order to obtain enough roots for trace elements analysis and equal amounts of media for root exudates extraction and metabolic profiling. After three weeks of growth, no visible phenotypic differences among WT and mutant plants could be highlighted: all genotypes grown in Fe-deficient medium (10 μM Fe^2+^) were paler and plants grown in Fe-depleted solution (0 μM Fe^2+^) were chlorotic (Figure S7). Interestingly, chlorophyll content was slightly higher in both mutant lines in all media tested, while chlorophyll a/b ratio was significantly higher in Col-0 plants in control media and slightly higher in Fe-deficient and Fe-depleted media (Figure S8). However, the targeted metabolite profiling of root extracts and root exudates showed that under Fe-depleted conditions both mutant lines accumulated lower levels of various coumarins (scopoletin, scopolin, umbelliferone, skimmin) (Figure 7A), but secreted significantly higher amounts of scopolin in comparison to WT plants (Figure 7B).

**Figure 7.**
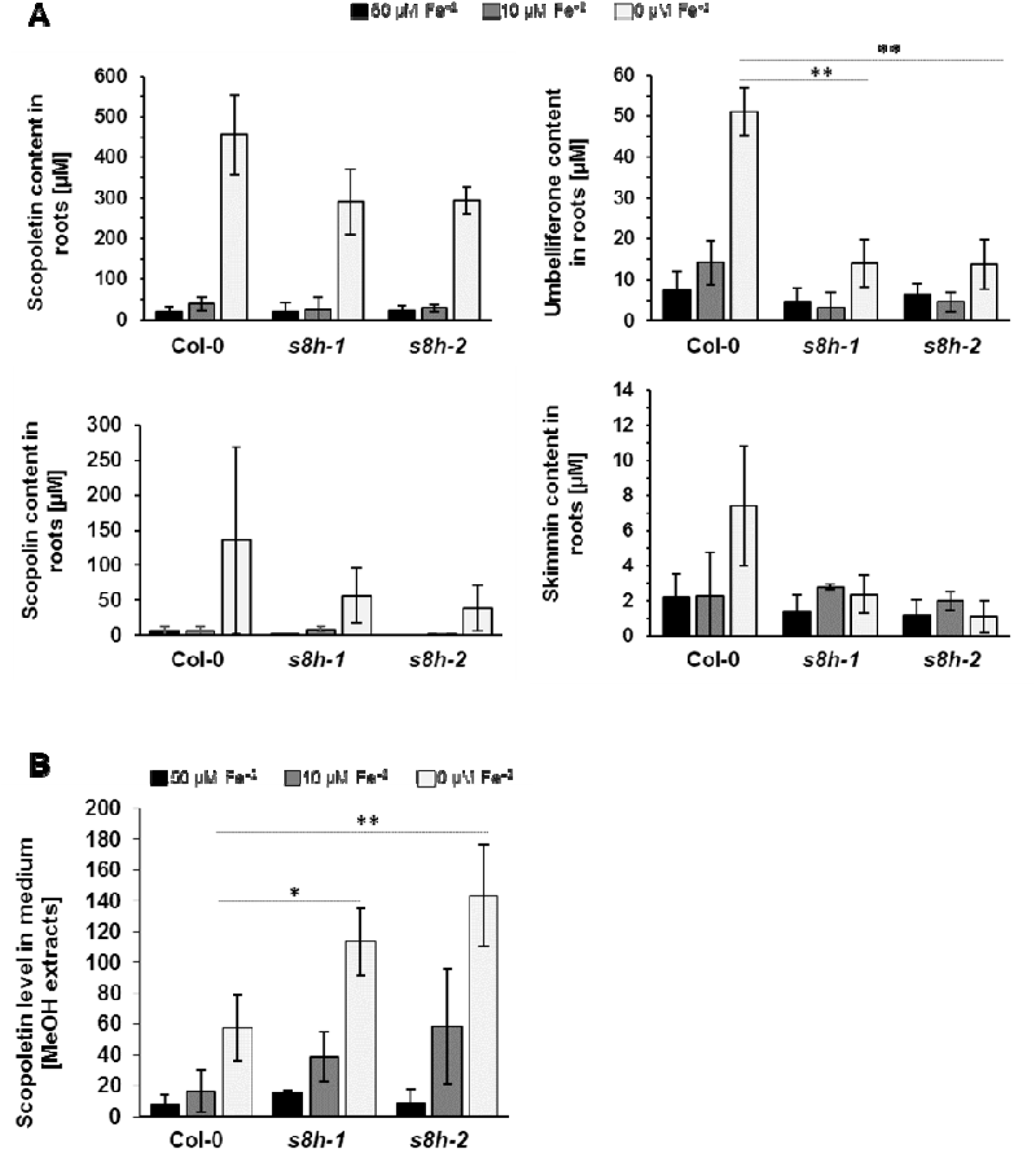
Biochemical characterization of *Arabidopsis thaliana s8h* mutants and Col-0 plants grown *in vitro* in 0.25 MS liquid cultures with various Fe content (0, 10 and 50 μM Fe^2+^). Relative levels of scopoletin, scopolin, umbelliferone and skimmin **(A)** in the methanol root extracts **(B)** and root exudates of *s8h* mutants and Col-0 plants. Metabolite profiling of coumarins were done by UHPLC. Error bars represent the SD from three measurements. (*) p < 0.05, (**) p < 0.01.

The trace element analysis of plants grown in liquid cultures media with various Fe levels indicated that the Fe content of both mutant lines (*s8h-1* and *s8h-2*) grown in Fe-depleted medium (0 μM Fe^2+^) were significantly lower (P < 0.05) in comparison to the corresponding control (Col-0) (Figure 8). Interestingly, the concentration of a range of other microelements heavy metals (Mn, Zn, Cu, Co, Cd) was also significantly modified in the *s8h* mutants compared to control plants (P < 0.05) grown in Fe-depleted medium (Figure 8).

**Figure 8.**
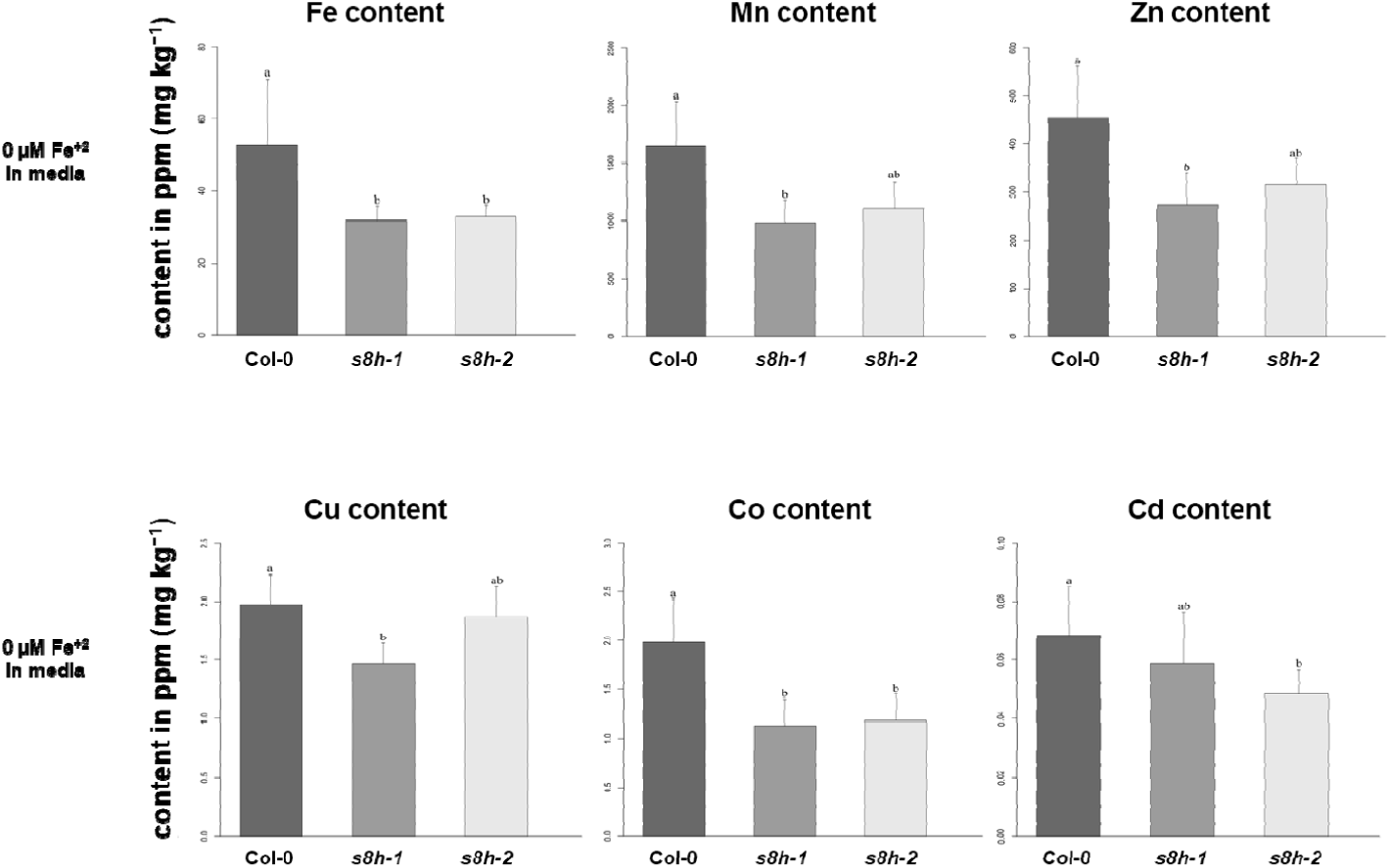
Microelements heavy metals content of Col-0 plants and *s8h* mutant roots grown *in vitro* for three weeks in Fe-depleted (0 μM Fe^2+^) 0.25 MS liquid culture. Microelement concentrations shown in ppm (mg kg^-1^) were determined by Inductively Coupled Plasma-Mass Spectrometry (ICP-MS). The results of one representative experimental replicate are presented. Error bars represent the SD from five biological replicates. Values that are not significantly different are indicated by the same letters. In all tests p-value < 0.05 was used.

In the light of recent results concerning regulatory pathways between phosphate (Pi) and Fe deficiency-induced coumarin secretion (Ziegler et al., 2016) and our soil experiment results in which Col-0 and *s8h-1* phenotypes were depending on the P to Fe ratio (Figure S9, Table S6), another interesting observation from the trace elements analysis is the clearly lower levels of P content in both mutants (significantly lower in *s8h-1* line, P < 0.05) in comparison to Col-0 plants under Fe-depleted (0 μM Fe^2+^) conditions (Figure S10A).

Plants grown in Fe-deficient (10 μM Fe^2+^) and Fe-optimal (50 μM Fe^2+^) media did not show clear changes in trace elements content (Figure S10ABC). The only clear exception was significantly higher Mo content in *s8h* mutants grown in Fe-deficient medium (Figure S10C).

### Phenotyping of s8h mutants grown on MS plates and in hydroponic solutions with various concentrations of Fe and other microelements

The impact of trace elements on Arabidopsis growth has been deeper investigated by cultivating the *s8h* mutants and control plants on plates and hydroponic cultures characterized by different Fe concentrations and other microelements content.

To be able to observe the growth of both rosettes and roots, the MS plates were placed in a vertical and horizontal position. Interestingly, the growth of all genotypes was better on 0.25 MS medium than on the corresponding plates with 0.5 MS, irrespective of the concentration of Fe ions (Figure S11). These differences were particularly striking after three weeks of cultivation. When grown on 0.25 MS Fe-deficient (10 μM Fe^2+^) medium, the *s8h* mutants showed a strong reduction in fresh weight (Figure S12A), were becoming much paler and had shorter and less lateral roots when compared to Col-0 plants (Figure 9A). On 0.5 MS Fe-deficient medium both genotypes display a significant reduction in shoots and roots growth, but *s8h* mutants showed a smaller rosettes size than Col-0 plants (Figure 9B). Under both Fe-depleted conditions (0 μM Fe^2+^), mutant lines and Col-0 plants showed significantly smaller and chlorotic shoots while roots were greatly shorter (Figure 9A-B). However, the wild type plants grew slightly better. Interestingly, the shoots of *s8h* mutants grown on 0.25 MS Fe-optimal (50 μM Fe^2+^) medium were clearly larger in comparison to the WT plants, which was not the case on 0.5 MS medium with optimal Fe content (Figure 9A-B).

**Figure 9.**
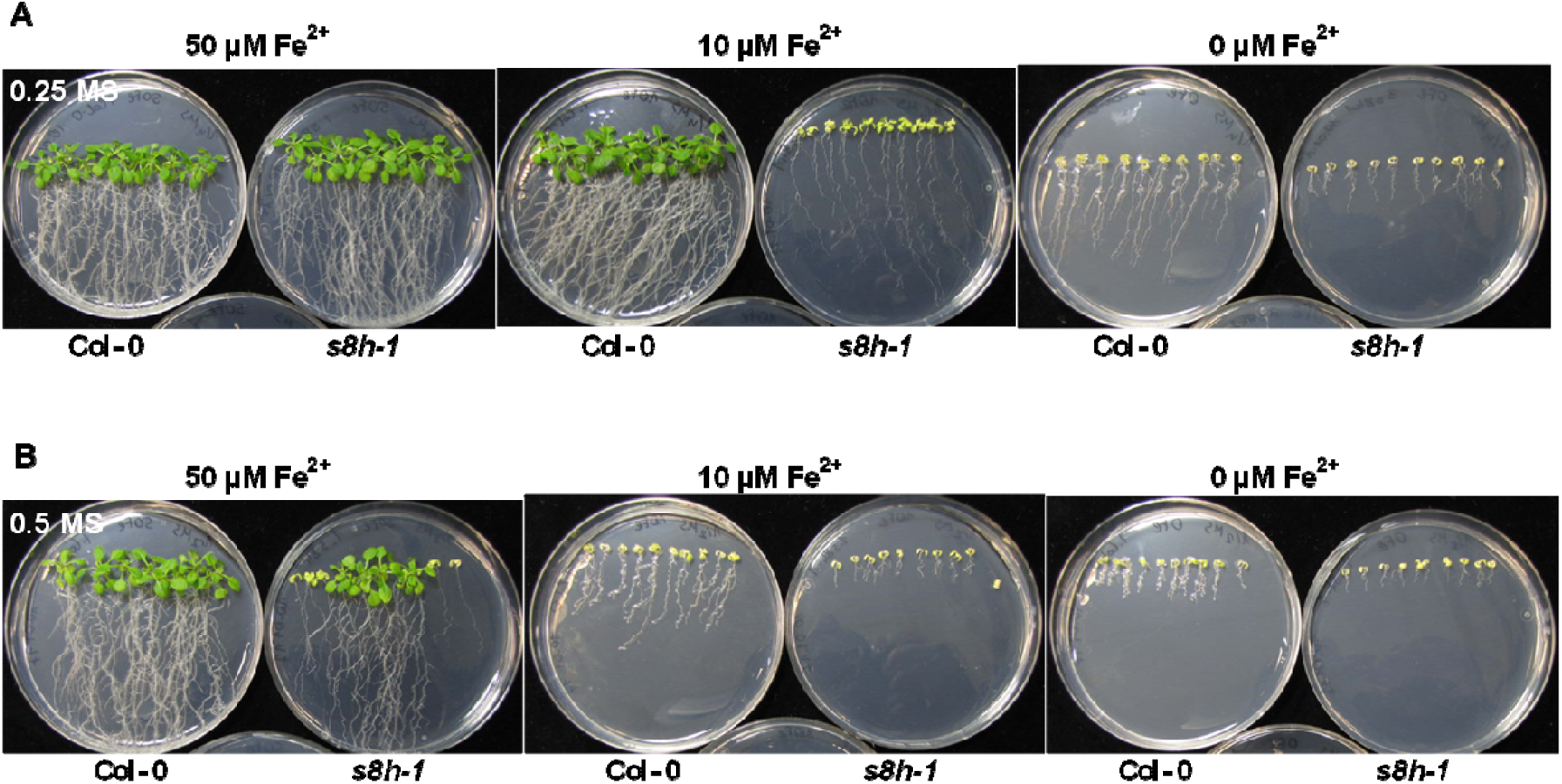
Phenotypic appearance of three weeks old Col-0 and *s8h-1* plants grown on (A)0.25 MS and (B) 0.5 MS media. Plants were grown Murashige and Skoog’s (MS) media with various Fe availability (0-50 μM Fe^2+^) in plant growth chambers under a photoperiod of 16 h light (∼5000 lux) at 22 °C and 8 h dark at 20 °C. Due to the limited space, only *s8h-1* line is shown, a second mutant allele (*s8h-2*) showed a very similar respond.

Observation of plants grown on Fe-deficient plates (10 μM Fe^2+^) under UV light showed that both *s8h* mutant lines secreted to media an increased level of fluorescent compound (Figure S13A), which might be related to significantly higher amounts of scopoletin detected in *s8h* Fe-deficient liquid culture solution (Figure 7B). This difference of fluorescence could not be observed in control conditions (50 μM Fe^2+^), in which mutant plants seemed to accumulate slightly more fluorescent compounds in roots compared to WT plants (Figure S13B).

Similar phenotypic variation in plant responses to various Fe content depending on other micronutrients concentration we observed in hydroponic cultures, during which two types of modified Heeg media were used (Table S3 for details). This way, we made evidence that both *s8h* mutants when grown in Fe-depleted (0 μM Fe^2+^) solution with 10 times less microelements, were clearly paler than Col-0 plants (Figure S14C). This phenotype was associated with lower amounts of chlorophylls. Interestingly, both mutant lines under Fe-deficient (10 μM Fe^2+^) conditions were much larger when compared to WT plants with greatly increased fresh weight (Figure S14B). Surprisingly, when grown with optimal Fe (40 μM Fe^2+^) but lower amounts of other micronutrients, all genotypes were significantly smaller but did not show any chlorosis symptoms or changes in chlorophyll content (Figure S14A).

In contrast, when plants were cultivated in 10xHeeg solution characterised by a higher amount of micronutrients, there were no visible differences among *s8h* mutants and WT plants when grown in presence of different Fe level (Figure S15). Under optimal and Fe-deficient conditions, all plants display a WT-like phenotype with normal pigmentation, but in solution without Fe all plants were smaller and chlorotic. It should be highlighted that in the above described hydroponic experiments, both types of solutions (1xHeeg and 10xHeeg) were fully changed once per week.

Since the recent literature data provide support for a crucial role of root-secreted coumarins in the acquisition of Fe (Tsai and Schmidt, 2017), we decided to repeat the above hydroponic cultures but without changing nutrient solution once per week. Instead, boxes were refilled by the addition of a fresh medium to keep the similar volume of solution in each culture and not to get rid of the potentially secreted coumarins. When grown in solution with lower micronutrients content (1xHeeg), under Fe-depleted conditions (0 μM Fe^2+^) both *s8h* mutants were becoming chlorotic (Figure S16C) as previously observed (Figure S14C). But the growth of all genotypes was not clearly affected in optimal solution and under Fe-deficiency (40 and 10 μM Fe^2+^, respectively) (Figure S16AB), which could be due to a higher coumarin accumulation in solutions that were not fully changed. Similarly to previously conducted hydroponic experiments, all plants grown in 10xHeeg solution under Fe-deficient and optimal conditions did not show any sign of chlorosis or growth retardation (Figure S17AB). But surprisingly, under Fe-depleted conditions all genotypes were only slightly smaller when compared to optimal conditions and had normally pigmented leaves with no changes in chlorophyll content (Figure S17C). Only, shoots of mutant lines seemed to be slightly brighter.

## Discussion

In the Arabidopsis genome there are more than 100 genes encoding enzymes sharing homologies with dioxygenases. The 2OGD is the second largest enzyme family in plants whose members are involved in various oxygenation/hydroxylation reactions (Kawai et al., 2014), including biosynthesis of coumarins that are important compounds contributing to the adaptation of plants to biotic and abiotic stresses (Kai et al., 2008; Vialart et al., 2012; Matsumoto et al., 2012; Rodriguez-Celma et al., 2013a). Among the most common stress factors leading to plant growth disorders and chlorosis are micronutrients deficiency or excess that can result in various physiological disorders (Marschner, 2012). Maintaining the nutrient homeostasis in cells is crucial for the proper functioning of plants and the mechanisms governing minerals uptake and transport must be strictly controlled.

Plants use different strategies to compensate Fe limitation. Recently, it was shown that one of the Arabidopsis Fe(II)- and 2OG-dependent dioxygenase, the scopoletin synthase F6’H1 (Kai et al., 2008), is required for the biosynthesis of the Fe(III)-chelating coumarin esculetin that is released into the rhizosphere as part of the Fe uptake by Strategy I plants (Schmid et al., 2014).

Here, we have conducted the analysis of a strongly Fe-responsive gene At3g12900 of unknown biological function, which shares high homologies with the Fe(II)- and 2OG-dependent dioxygenase family, to possibly reveal its contribution to Fe homeostasis in plants. The At3g12900 gene was selected based on its high homology to earlier described F6’H1 dioxygenase which was proven to play a crucial role in Fe acquisition under alkaline soil conditions (Schmid et al., 2014), and the literature data on plant responses to Fe deficiency at the transcriptome and proteome level (Lan et al., 2011; Rodriguez-Celma et al., 2013b; Fourcroy et al., 2014; Schmidt et al., 2014). Similarly to F6’H1, the protein encoded by At3g12900 accumulates several folds in Fe-deficient roots in comparison to Fe-sufficient ones (Lan et al., 2011). By analysing large microarray datasets both genes (At3g12900 and At3g13610 encoding F6’H1) were found to be positively correlated with genes actively involved in Fe deficiency response (Vigani et al., 2013) such as *IRT1*, *FRO2*, *CYP82C4* (Murgia et al., 2011), *Ferroportin/Iron-Regulated* (*IREG2*) (Morrissey et al., 2009) and encoding metal tolerance protein (*MTP3*; Arrivault et al., 2006). Moreover, according to the expression data present at TAIR the At3g12900 is expressed specifically in roots. The root tissue is the site of coumarin accumulation induced in response to various environmental stresses including Fe limitation (Schmid et al., 2014).

We determined the substrate specificity of enzyme encoded by At3g12900 to test its possible involvement in coumarin biosynthesis as an important part of the Fe uptake strategy in Arabidopsis. An *in vitro* enzymatic activity assay revealed that this enzyme is involved in the conversion of scopoletin into fraxetin *via* hydroxylation at the C8-position and was named S8H. The Michaelis constant (*K*_m_= 11 μM) determined in our *in vitro* experiments was in the average similar to those reported for the biosynthetic enzymes of specialized metabolism (Kai et al., 2008; Vialart et al., 2012). These results taken together suggest that the hydroxylation of scopoletin (and as an effect the synthesis of fraxetin) is the main activity of the S8H enzyme. The experiments performed with *in vitro-*produced enzymes also made evidence that the S8H activity was dependent on the concentration of Fe^2+^.

Transient expression of the protein in *N. benthamiana* plants and a subsequent metabolic analysis done on the infiltrated leaves further confirmed the results of *in vitro* assay. A simultaneous transformation of *S8H* and *F6’H2* heterologous genes in tobacco leaves resulted in a significantly higher accumulation of fraxetin; this concentration was intermediate when *F6’H2* alone was expressed and much lower when empty vector or *S8H* alone were expressed. The latter one could be explained by the fact that tobacco plants do not synthesize scopoletin in a constitutive way but synthetize fraxetin. An overproduction of F6’H2 therefore induce the synthesis of scopoletin from feruloyl-CoA, which is naturally present in tobacco leaves (Kai et al., 2008), and provide this way the substrate for the reaction catalysed by S8H resulting in significantly higher content of fraxetin.

To better understand the link between Fe-homeostasis in plants and the biosynthesis of fraxetin, and consequently to show that the *in vitro* enzyme activity of S8H is relevant *in vivo,* we performed a detailed phenotypic characterization of Col-0 plants and two independent *s8h* mutant lines grown under different Fe regimes using various types of culture. Plants were grown in hydroponic solution, soil mixes, *in vitro* liquid cultures and on MS plates with various Fe and other micronutrients availability. Our results clearly showed that the *s8h* plants carrying mutated S8H alleles are strongly affected by Fe-deficient conditions. Targeted metabolite profiling of *s8h* mutants demonstrated that coumarin profiles are significantly modified in mutant roots grown in Fe-depleted conditions. We detected higher amounts of scopoletin in exudates from *s8h* mutant roots grown in liquid cultures. It was associated with lower levels of various coumarins and lower Fe content in mutants roots as compared to the wild-type plants. The *s8h* rosettes grown in 1xHeeg Fe-depleted hydroponic solutions were clearly paler than Col-0 plants. It was associated with striking changes in metabolite profiles of coumarins in mutant roots. In comparison to Col-0 plants, under this condition we detected a significantly higher accumulation of scopoletin and scopolin in *s8h* roots that can suggest the inhibition of scopoletin-hydroxylation-dependent synthesis of fraxetin in mutant tissues. Most likely, *in vitro* culturing conditions in a small volume of liquid solution favor increased secretion of exudates and therefore we observed lower levels of various coumarins in *s8h* roots grown in liquid cultures linked with a significantly higher scopoletin content in mutant roots exudates. The phenotypic differences between genotypes were also apparent on MS plates containing Fe-deficient medium, on which the *s8h* mutants showed chlorosis, significant growth retardation and secreted an increased level of fluorescent compounds compared to WT plants.

Taking into account the results of soil experiments in which plants were cultured in soil mixtures with various chemical composition, it will be also interesting to test the growth and metabolic profiles of roots and root exudes of *s8h* mutants grown in hydroponic solutions with various P availability. The fact that *s8h-1* mutants were larger in soil mix characterized by a relatively low level of P and high level of Fe, is particularly interesting in the light of recent reports on the common and antagonistic regulatory pathways between phosphate (Pi) and Fe deficiency-induced coumarin secretion (Ziegler et al., 2016).

Another interesting aspect for further investigation is linked to the ICP-MS results indicating that additionally to Fe a range of other heavy metals (Mn, Zn, Cu, Co, Cd) were significantly decreased in the *s8h* mutants, and to the phenotypic variation we observed in hydroponic cultures and on MS plates depending on both Fe and other micronutrient contents. The latter one could be expected due to the well-known phenomenon of interdependence of individual micronutrients from each other (Ihnatowicz et al. 2014) and the fact that various heavy metals interfere with Fe-deficiency responses (Leskova et al. 2017). Nevertheless, we proved that to get a broad overview of plant responses to nutrient deficiencies and to better understand the physiological role of involved genes/enzymes one need to consider using various types of cultures, solution and media in experimental procedure.

Among numerous targeted metabolite profiling experiments conducted, we clearly and repetitively obtained results showing significantly changed coumarin profiles in both *s8h* mutant lines. We observed a significantly higher content of scopoletin in the root tissue in hydroponic experiments and in the root exudates in liquid cultures. Scopoletin is a substrate for the reaction catalysed by S8H. The lack of scopoletin 8-hydroxylase in the *s8h* mutant background could lead to higher levels of scopoletin and its corresponding glycoside scopolin.

At the moment, we have no explanation for the fact that only small amounts of fraxetin were detected in some of the Col-0 samples under Fe-deficiency conditions (data not shown). It should also be mentioned that unexpectedly in one experimental replicate we detected relatively high amounts of fraxin and a low level of fraxetin in *s8h-1* mutant background, as well as low amounts of fraxin in some other Col-0 and *s8h* replicates. It cannot be excluded that the above described queries could be explained by the presence of another enzyme involved in fraxetin biosynthesis or alternative metabolic pathway being induced in *s8h* mutant background. The synthetized fraxetin in Col-0 plants could also be further demethylated to 6,7,8-trihydroxycoumarin with beneficial effect for plants under Fe-deficiency or alternatively fraxetin could be directly involved in increasing Fe availability. The latter one cannot be excluded taking into account a catechol-type structure of fraxetin.

This needs to be further investigated. As presented by Schmid et al. (2014), Fe-deficient chlorotic phenotype of *f6’h1* seedlings grown under low Fe availability could be reversed by exogenous application of esculetin, esculin and scopoletin. In parallel, the result of *in vitro* assay showed that only esculetin was able to chelate and mobilize Fe^3+^ (Schmid et al., 2014). This suggests that compounds bearing an ortho-catechol moiety, such as esculetin and fraxetin, may be involved in coumarin secretion for Fe acquisition. The beneficial effect of exogenous application of scopoletin on the reversion of *f6’h1* chlorotic phenotype demonstrated by Schmid et al. (2014), could be due to the activity of S8H enzyme catalyzing hydroxylation of scopoletin leading to fraxetin formation. Whatever mechanisms underlies such plant responses, given the results of *in vitro* enzyme activity and significant changes in metabolite profiles of coumarin metabolism in *s8h* mutants grown in Fe-depleted condition, it seems evident that the At3g12900 oxidoreductase is scopoletin 8-hydroxylase involved in fraxetin biosynthesis.

The presented results indicate that At3g12900 is an important candidate for playing essential role in Fe acquisition by plants. Fraxetin together with other root-released phenolic compounds, mainly esculetin deriving from scopoletin and scopolin, have been suggested to be Fe chelators or Fe uptake facilitators in Arabidopsis (Schmid et al., 2014; Fourcroy et al., 2014; Schmidt et al., 2014; Brumbarova et al., 2015) and that esculetin and fraxetin or fraxetin derived compounds are possibly involved in the transport of Fe ions into the plant cell under Fe deficiency. The precise physiological function of the phenolic compounds synthesized by plants under Fe deficiency stress and the possible mechanism underlying plant responses to Fe limitation under calcareous conditions remains unknown (Schmid et al., 2014). Elucidating the biological role of the scopoletin 8-hydroxylase involved in coumarin biosynthesis is a prerequisite to the understanding of fraxetin function in Fe acquisition.

## Acknowledgments

We would like to thank Charles Clément for valuable technical assistance and Sabina Żołędowska for assistance in statistical analysis using by R version 3.3.2. This work was supported by the Polish National Science Centre grants: UMO-2014/15/B/NZ2/01073 and UMO-2014/12/T/NZ8/00316, and the LiSMIDoS PhD fellowship (UDA-POKL.04.01.01-00-017/1000).

## Additional information

Supplementary data are available in the online version of the paper.

**Figures S1.** Schematic representation of independent T-DNA mutant lines for At3g12900 gene.

**Figure S2.** The lack of *S8H* transcript in the *s8h-1* T-DNA mutant line confirmed by qRT-PCR.

**Figure S3.** Multiple-sequence alignment of amino acid sequences of F6’H1, F6’H2, S8H and ANS enzymes.

**Figure S4.** Results of recombinant his-tagged S8H protein purification.

**Figure S5.** Characterization of *Arabidopsis thaliana* S8H enzyme activity.

**Figure S6.** Kinetic parameters Vmax and *Km* of S8H.

**Figure S7.** Phenotypic appearance of Col-0 and *s8h* plants grown *in vitro* in liquid cultures with various levels of Fe.

**Figure S8.** Chlorophyll content and chlorophyll a/b ratio of Col-0 and *s8h* plants grown *in vitro* in liquid cultures with various levels of Fe.

**Figure S9.** Phenotyping and biochemical characterization of Col-0 and *s8h-1* plants grown in soil mixes with different Fe availability.

**Figure S10.** Trace element content of Col-0 and *s8h* plant roots grown *in vitro* in Fe-depleted liquid culture.

**Figure S11.** Phenotypic appearance of two weeks old Col-0 and s*8h-1* plants grown on (A)0.25 MS and (B) 0.5 MS media.

**Figure S12.** Fresh weight of three weeks old Col-0 and *s8h* plants grown *in vitro* on (A) 0.25 MS and (B) 0.5 MS media.

**Figure S13.** Phenotypic appearance of Col-0 and s*8h* plants observed under UV light.

**Figure S14.** Phenotypic characterization of Col-0 and *s8h* plants grown in 1xHeeg hydroponic solution (fully changed once per week) with various Fe content (0-40 μM Fe^2+^).

**Figure S15**. Phenotypic characterization of Col-0 and *s8h* plants grown in 10xHeeg hydroponic solution (fully changed once per week) with various Fe content (0-40 μM Fe^2+^).

**Figure S16.** Phenotypic characterization of Col-0 and *s8h* plants grown in 1xHeeg hydroponic solution (refilled with a fresh medium) with various Fe content (0-40 μM Fe^2+^).

**Figure S17.** Phenotypic characterization of Col-0 and *s8h* plants grown in 10xHeeg hydroponic solution (refilled with a fresh medium) with various Fe content (0-40 μM Fe^2+^).

**Table S1.** Genotyping of *s8h* mutant lines.

**Table S2.** A lack of *S8H* transcript in the *s8h* mutant backgrounds.

**Table S3.** Modified Heeg solutions used in hydroponic cultures.

**Table S4.** PCR amplification of the *S8H* ORF.

**Table S5.** Compounds used in the analysis of *in vitro* substrate specificity of S8H enzyme.

**Table S6.** Chemical analysis of soil mixes originated from different batches used in two experimental replicates.

